# Loss of EPAC2 results in altered synaptic composition of dendritic spines and excitatory and inhibitory balance

**DOI:** 10.1101/526772

**Authors:** Kelly A. Jones, Michiko Sumyia, Kevin M. Woolfrey, Deepak P. Srivastava, Peter Penzes

## Abstract

EPAC2 is a guanine nucleotide exchange factor that regulates GTPase activity of the small GTPase Rap and Ras) and is highly enriched at synapses. Activation of EPAC2 has been shown to induce dendritic spine shrinkage and increase spine motility, effects that are necessary for synaptic plasticity. These morphological effects are dysregulated by rare mutations of *EPAC2* associated with autism spectrum disorders. In addition, EPAC2 destabilizes synapses through the removal of synaptic GluA2/3-containing AMPA receptors. Previous work has shown that *Epac2* knockout mice *(Epac2^−/−^)* display abnormal social interactions, as well as gross disorganization of the frontal cortex and abnormal spine motility *in vivo.* In this study we sought to further understand the cellular consequences of knocking out *Epac2* on the development of neuronal and synaptic structure and organization of cortical neurons. Using primary cortical neurons generated from *Epac2^+/+^ or Epac2^−/−^* mice, we confirm that EPAC2 is required for cAMP-dependent spine shrinkage. Neurons from *Epac2^−/−^* mice also displayed increased synaptic expression of GluA2/3-containing AMPA receptors, as well as of the adhesion protein N-cadherin. Intriguingly, analysis of excitatory and inhibitory synaptic proteins revealed that loss of EPAC2 resulted in altered of expression of vesicular glutamate transporter 1 (VGluT1) and vesicular GABA transporter (VGAT), indicating a potential imbalance in excitatory/inhibitory inputs onto neurons. Finally, examination of cortical neurons located within the anterior cingulate cortex further revealed subtle deficits in the establishment of dendritic arborization *in vivo.* These data provide evidence that EPAC2 is required for the correct composition of synapses and that loss of this protein could result in an imbalance of excitatory and inhibitory synapses.

## Introduction

The ubiquitous second messenger molecule cyclic AMP (cAMP) is an important member of many signaling cascades in the central nervous system. cAMP signaling has been shown to be crucial for neuronal development, dendritic and axonal morphogenesis, and synaptic plasticity, and it modulates a broad range of cognitive functions, including working and reference memory (Lee, 2015; Ricciarelli and Fedele, 2018; Silva and Murphy, 1999). Alterations in upstream and downstream components of the cAMP pathway have also been shown to affect behaviors including sociability and communication (Burgdorf et al., 2007; Fischer and Hammerschmidt, 2011; Wang et al., 2008). Conversely, abnormal cAMP signaling has been implicated in a range of neurodevelopmental and psychiatric disorders, several of which affect cognitive functions (Garcia et al., 2016; Havekes et al., 2015; Kelley et al., 2008; Kelly et al., 2009; Nestler et al., 2002; Ricciarelli and Fedele, 2018).

cAMP signaling occurs via two main downstream pathways, one that is protein kinase A (PKA)-dependent and another that is PKA-independent (Bos, 2003). PKA-independent cAMP targets include EPAC (exchange protein directly activated by cAMP) proteins (Bos, 2003) and cyclic nucleotide-gated channels. While much attention has been dedicated to the role of the PKA-dependent pathway in plasticity and cognitive behavior, relatively little is known about the roles of the PKA-independent mechanisms in the brain. EPAC2, also known as cAMP-GEFII or RapGEF4, is a brain-enriched guanine-nucleotide exchange factor (GEF) for the small GTPase Rap and is the major EPAC protein expressed throughout development and in the adult brain (Kawasaki et al., 1998; Ulucan et al., 2007; Woolfrey et al., 2009). EPAC2 contains two cAMP-binding domains and a Rap-GEF domain, in addition to other domains. Binding of cAMP to the cAMP-binding domain enhances the catalytic activity of the GEF domain toward Rap in both EPAC1 and EPAC2 (Bos, 2003; Woolfrey et al., 2009). Work from our group has also shown that EPAC2 is required for the establishment and maintenance of basal dendritic arborization through its interaction with the small GTPase Ras during development (Srivastava et al., 2012b). Activation of EPAC2 in neurons with a mature cellular morphology results in the shrinkage of dendritic spines and synapse destabilization through the removal of GluA2/3-containg AMPA receptors from synapses (Woolfrey et al., 2009). Moreover, EPAC2 is a critical mediator of dopamine D1 receptor-mediated spine remodeling (Woolfrey et al., 2009). Interestingly, EPAC2 activation can also be regulated by the adhesion protein neuroligin 3 (NL3), a protein associated with autism spectrum disorders (ASDs) (Woolfrey et al., 2009). Critically, rare coding variants of *EPAC2* have also been associated with ASDs (Bacchelli et al., 2003), and these variants alter the ability of EPAC2 to regulate synaptic structure and function (Woolfrey et al., 2009). Interestingly, *Epac2* knockout mice *(Epac2^−/−^)* displayed abnormal organization of the anterior cingulate cortex (ACC), reduced spine dynamics *in vivo* (Srivastava et al., 2012a; Viggiano et al., 2015) and specific deficits in social and communicative behaviors (Srivastava et al., 2012a). These behavioral deficits are also mirrored in mice lacking both *Epac1* and *Epac2* (Yang et al., 2012; Zhou et al., 2016). While these data indicate a role for EPAC2 in both developing and adult brain, a comprehensive examination of this protein’s role in synaptic organization *in vitro* and *in vivo* has yet to be performed.

In this study, we have used primary cortical cultures generated from *Epac2^−/−^* mice and wild-type littermates (Srivastava et al., 2012a) to examine the ability of cells to respond to cAMP stimulation. Furthermore, we have examined the impact of EPAC2 loss on the organization of synapses on cortical neurons. Specifically, we have focused on the synaptic presence of AMPA receptors and adhesion proteins known to directly or indirectly be associated with EPAC2. We further investigate whether loss of EPAC2 altered the ratio of excitatory and inhibitory inputs on neurons. Finally, as we have previously shown that loss of *Epac2* alters the dendritic organization and spine dynamics of layer 2/3 and layer 5 cortical neurons, respectively, located in pre-motor and somatosensory areas (Srivastava et al., 2012a; Srivastava et al., 2012b), we examined whether knockout *Epac2* alters the dendritic and synaptic morphology of layer 5 neurons located in the ACC. The result of these investigations indicate loss of EPAC2 impacts the abundance of AMPA receptor subunits and specific adhesion proteins at synapses. Moreover, *Epac2^−/−^* neurons display altered excitatory and inhibitory inputs. Finally, layer 5 ACC neurons display subtle alterations in dendritic arborization in *Epac2^−/−^* mice. These data indicate that EPAC2 is required for the normal organization of the synaptic proteome during development, as well as balanced excitatory and inhibitory input to cortical neurons.

## Materials and Methods

### Reagents

cAMP analog 8-(4-chloro-phenylthio)-2’-O-methyladenosine-3’,5’-cyclic monophosphate (8-CPT) was purchased from Tocris Bioscience (R&D Systems). Sources of antibodies are as follows: rabbit anti-EPAC2 polyclonal (Cell Signaling Technology), mouse anti-βactin monoclonal (Sigma), rabbit anti-NL3 polyclonal (Santa Cruz Biotechnology), rabbit anti-GluA2/3 polyclonal (Millipore), rabbit anti-VGAT polyclonal (Millipore), mouse anti-VGluT1 monoclonal (Millipore), mouse anti-PSD-95 monoclonal clone K28/43 (University of California-Davis/National Institutes of Health Neuromab Facility), chicken anti-GFP polyclonal (Abcam) and mouse anti-bassoon monoclonal (Abcam).

*Epac2^−/−^* mice (C57BL/6) were generated as previously described (Srivastava et al., 2011). In order to label a subset of layer 5 neurons with green fluorescent protein (GFP), wild-type and *Epac2^−/−^* mice were crossed with the Tg(*Thy1*-GFPM)2Jrs/J transgenic line (Jackson Labs) as previously described (Srivastava et al., 2012a; Viggiano et al., 2015). Tg(Thy1-GFPM)2Jrs/J express GFP in a subset of layer 5 neurons throughout the neocortex; labelled neurons are ideal for morphological studies. This resulted in the generation of Epac2^+/+GFP^ and *Epac2^-/-GFP^* mice. Mice were used in accordance with ACUC institutional and national guidelines under approved protocols.

### Culturing of primary cortical neurons from wild-type and Epac2^−/−^ mice

Dissociated cultures of primary cortical neurons were prepared from *Epac2* wild-type and knockout mice. The latter were created by Professor Susumu Seino of Kobe University (Shibasaki et al., 2007). Cortical neuronal cultures, consisting of mixed sexes, were prepared from P0 mouse pup in accordance with ACUC institutional and national guidelines under approved protocols and as described before (Srivastava et al., 2012a; Srivastava et al., 2011). Briefly, mouse pups were euthanized by decapitation, brains were quickly removed, and cortical tissue was isolated, digested, and dissociated. Cells were plated onto 18 mm glass coverslips (No 1.5; 0117580, Marienfeld-Superior GmbH & Co.), coated with poly-D-lysine (0.2mg/ml, Sigma), at a density of 3−10^5^/well equal to 857/mm^2^. Neurons were cultured in feeding media: neurobasal medium (21103049) supplemented with 2% B27 (17504044), 0.5 mM glutamine (25030024) and 1% penicillin:streptomycin (15070063) (all reagents from Life technologies). Neuron cultures were maintained in presence of 200 μM D,L-amino-phosphonovalerate (D,L-APV, ab120004, Abcam) beginning on DIV (days *in vitro)* 4 in order to maintain neuronal health for long-term culturing and to reduce cell death due to excessive Ca^2+^ cytotoxicity via over-active NMDA receptors (Srivastava et al., 2011). Half media changes were performed twice weekly until desired age (DIV 23-25). A subset of primary cortical neurons were transfected with eGFP or control or Epac2-shRNA at DIV 21 for 2 or 5 days respectively, using Lipofectamine 2000 (11668027, Life Technologies) (Srivastava et al., 2011). Briefly, 2-4 μg of plasmid DNA was mixed with Lipofectamine 2000 and incubated for 4-12 hours, before being replaced with fresh feeding media. Transfections were allowed to proceed for 2 days after which cells were used for pharmacological treatment or immunocytochemistry (ICC).

### Immunocytochemistry (ICC)

Neurons were washed in PBS and then fixed in 4% formaldehyde/4% sucrose PBS for 10 minutes at room temperature followed by incubation in methanol pre-chilled to -20°C for 10 minutes at 4°C. Fixed neurons were then permeabilized and blocked simultaneously (2% Normal Goat Serum, 5425S, New England Biolabs and 0.1% Triton X-100) before incubation in primary antibodies overnight and subsequent incubation with secondary antibodies the following day (Srivastava et al., 2011).

### Quantitative Analysis of Spine Morphologies and Immunofluorescence

Confocal images of double-stained neurons were acquired with a Zeiss LSM5 Pascal confocal microscope and a 636 objective (NA = 1.4). Two-dimensional maximum projection images were reconstructed and analysed using MetaMorph software (Molecular Devices, Sunnyvale, CA, USA) (Srivastava et al., 2011). Morphometric analysis was performed on spines from two dendrites (secondary or tertiary branches), totaling 100 μm, from each neuron. Linear density (per 10 μm) and spine area, length and breadth was measured automatically using MetaMorph Software (Molecular Devices) (Srivastava et al., 2011). Protein clustering was imaged as above. Resultant images were background-subtracted and thresholded equally to include clusters with intensity at least 2-fold above the adjacent dendrite. Analyses of puncta were performed on spines from at least two dendrites (secondary or tertiary branches), totaling 100 μm, from each neuron. The linear density (number per 10 μm of dendrite length) and total gray value (total immunofluorescence intensity) of each synaptic protein cluster was measured automatically using MetaMorph (Srivastava et al., 2011). Co-localized puncta were defined as puncta that contained immunofluorescence staining greater than background of the reciprocal protein costained; background fluorescence was the average background intensity from five regions of interest plus two standard deviations (Glynn and McAllister, 2006). Cultures that were directly compared were stained simultaneously and imaged with the same acquisition parameters. For each condition, 10–16 neurons from at least 3 separate experiments were used. Experiments were conducted blind to condition and on sister cultures. In the green/magenta colour scheme, co-localization is indicated by white overlap.

### Western blotting

Whole cell lysates were prepared from DIV 25 neurons generated from wildtype or knockout mice. Cells were lysed in RIPA buffer (150 mM NaCl, 10 mM Tris-HCl (pH 7.2), 5 mM EDTA, 0.1% SDS (wt/vol), 1% Triton X-100 (vol/vol), 1% deoxycholate (wt/vol), and inhibitors), before being sonicated with 10 short bursts. Sample buffer was added to all samples, which were then denatured for 5 minutes at 95°C and stored at -80°C until used further. All samples were subsequently separated by SDS-PAGE and analyzed by Western Blotting with antibodies against EPAC2 and β-actin. Quantification of bands was performed by measuring the integrated intensity of each band and normalizing to β-actin, for protein loading, using Image J.

### Preparation of cortical tissue sections

In order to examine dendritic and synaptic structures in cortical layer 5, Epac2^+/+GFP^ and *Epac2*^−/−,GFP^ mice were anesthetized with a ketamine/xylazine mixture and perfused transcardially with PBS followed by 4% paraformaldehyde in PBS 8 weeks of age. All experiments were carried out in accordance with ACUC institutional and national guidelines under approved protocols. Brains were removed, postfixed overnight in 4% paraformadehyde/PBS, and cryoprotected in 30% sucrose/PBS. Brains were then embedded in 3% agarose and sectioned coronally at 300 μm with a vibratome. Sections were mounted onto a glass slide and covered with a No.1.5 glass coverslip with 2 #1 coverslips (~150 μm thickness) placed either side of the section to avoid damage to the tissue (Srivastava et al., 2012b).

### 2 photon laser scanning microscopy (2PLSM) imaging of fixed brain sections and *quantitative* morphological analysis

Fixed brain sections were imaged on a Olympus BX51-WIF upright, fixed-stage microscope using a Zeiss LD LCi PA 25x/0.8NA multi-immersion lens (440842-9870–000000), and a Coherent Chameleon-Ultra2 tunable (680 nm to 1080 nm) laser system utilizing Ti:sapphire, attenuated by two ConOptics Pockels cell electro-optic modulators. The scanning system software used was *LaserSharp* (BioRad). GFP-expressing cells were excited at 950 nm, and Z-stacks (100-200 images) were acquired at 500 lines per second, at a resolution of 1024 x 1024 pixels: at digital zoom = 1 (for dendritic arbors), xy pixel = 271 0.45um with 1 um Z steps; at digital zoom = 3.6 (for dendritic spines), xy pixel = 0.13 um with 0.75um Z-steps. Kalman corrections (N=6) were applied to images acquired at zoom of 3.6. The ACC was identified and only GFP-expressing layer 5 pyramidal neurons in the ACC were imaged. Only cells exhibiting intact healthy secondary and tertiary apical and basal dendrites were imaged and used for quantification. Following acquisition, images were projected as 2-D Z-projections using MetaMorph for analysis of dendritic spines. For each condition, 1-2 cells from 6-8 animals imaged. Two dendrites between 50 and 100 μm in length per cell were measured: only spines on tertiary apical or secondary basal dendrites were imaged to reduce variability. Dendritic spine density (number of spines per 10 μm) as well as spine area, length and breadth were calculated using Metamorph. To examine dendritic arborization, z-stacks were maintained in 3 dimensions during tracing of dendritic arbors using the Neuromantic program (http://www.reading.ac.uk/neuromantic/), (Myatt et al., 2012). Briefly, neurites were digitally traced and subsequently reconstructed in 3D using Neuromantic. SWC data files, encoding the 3D reconstruction of the dendritic arbors, were exported and analyzed using L-measure (Scorcioni et al., 2008).

### Statistical Analysis

All statistical analysis was performed in GraphPad. Differences in quantitative immunofluorescence, dendritic spine number were probed by one-way-ANOVAs with Tukey correction for multiple comparisons. Error bars represent standard errors uless stated otherwise.

## Results

### Epac2^−/−^ neurons exhibit abnormal dendritic spine morphology in response to 8-CPT stimulation

Previous studies have demonstrated a role for an EPAC2-dependent regulation of dendritic spine morphology in response to cAMP stimulation (Woolfrey et al., 2009). Thus, we were interested to see if primary cortical neurons (days *in vitro* [DIV] 21-23) from wild-type and *Epac2* knockout mice differed in their ability to respond to cAMP. Western blotting of cell lysates from primary cultures confirmed loss of EPAC2 in knockout cultures (**Figure 1A**). Next, we compared the dendrite density and size of dendritic spines in neurons from wild-type and *Epac2r^−/−^* mice and found no difference in density, but an increase in spine area in neurons from knockout mice (**Figure 1B-C**). When we stimulated neurons from wild-type and *Epac2r^−/−^* mice with 8-CPT to mimic a PKA-independent cAMP signaling mechanism, analysis of spine morphology revealed that 8-CPT caused shrinkage of dendritic spines in neurons from wild-type cultures but not *Epac2r^−/−^* cultures (**Figure 1D-F**). 8-CPT treatment caused no differences in spine density in either wild-type or knockout neurons. Taken together, these data indicate that knockout of *Epac2* abolishes cAMP-dependent regulation of dendritic spine morphology.

**Figure 1.**
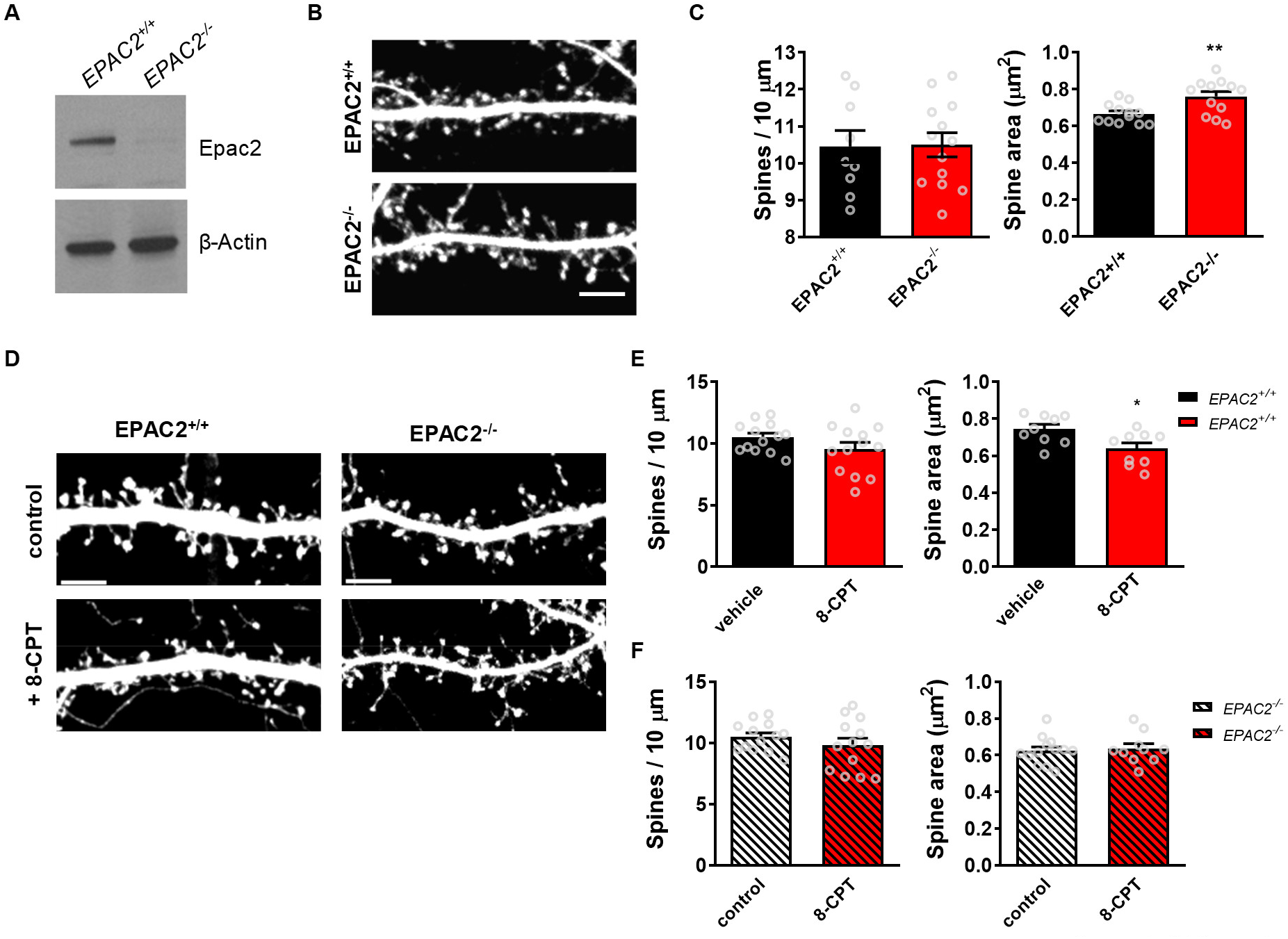
Dendritic spine morphology in pyramidal neurons from cortical cultures from wild-type or *Epac2^−/−^* mice. **(A)** Western blot of whole cell lysates from wild-type (Epac2^+/+^) and *Epac2^−/−^* cortical cultures demonstrating loss of EPAC2 in knockout cells. **(B)** Representative confocal images of dendritic spines on cortical neurons (DIV 24) from wild-type or *Epac2^−/−^* cultures. Scale bar=5 μm. **(C)** Quantification of spine linear density and area from the images in panel B. **P=0.0040, Student’s t test; n=12-17 cells from 3-4 independent cultures/genotype. (D) Confocal images of dendritic spines on cortical neurons from wild-type or *Epac2^−/−^* mice, treated with vehicle control or 8-CPT. Scale bars=5 μm. **(E)** Quantification of effect of 8-CPT treatment on spine linear density and area in wild-type neurons. *P=0.0183, Student’s t test; n=10-13 cells per condition, 4 independent cultures/genotype. **(F)** Quantification of effect of 8-CPT treatment on spine density and area in *Epac2^−/−^* neurons. P=0.8491, Student’s t test; n=11-13 cells per condition, 4 independent cultures/genotype. Data are presented as means ± s.e.m.; each data point represents an individual cell.

### Epac2^−/−^ neurons have increased synaptic levels of GluA2/3

We have previously shown that EPAC2 interacts with other postsynaptic proteins, including PSD-95 and GluA2 (Woolfrey et al., 2009). Moreover, EPAC2 regulates the trafficking of GluA2/3-containing glutamate receptors and AMPA receptor-mediated transmission (Woolfrey et al., 2009). To examine whether loss of EPAC2 altered the synaptic content of AMPA receptors, we generated primary cultures of cortical neurons from wild-type and *Epac2r^−/−^* mice and immunostained them for the synaptic protein PSD-95 and GluA2/3-containing AMPA receptors. When we examined PSD-95 puncta density, we found no differences between the genotypes (**Figure 2A-B**), indicating that loss of *Epac2* did not affect the density of synapses. This is consistent with the observation that *Epac2^−/−^* neurons do not have altered spine number. However, when we examined GluA2/3 puncta density, we found a significant increase in neurons from *Epac2^−/−^* mice compared to wild-type mice (**Figure 2A, C**). Furthermore, GluA2/3 clusters were larger in *Epac2r^−/−^* cultures (**Figure 2A, D**). These effects on puncta size and density were also accompanied by an increase in the number of PSD-95 and GluA2/3 colocalized puncta in *Epac2r^−/−^* neurons (**Figure 2A, E**), indicating an increased presence of GluA2/3 at synapses. These data further demonstrate that EPAC2 regulates the synaptic content of GluA2/3-containing AMPA receptors.

**Figure 2.**
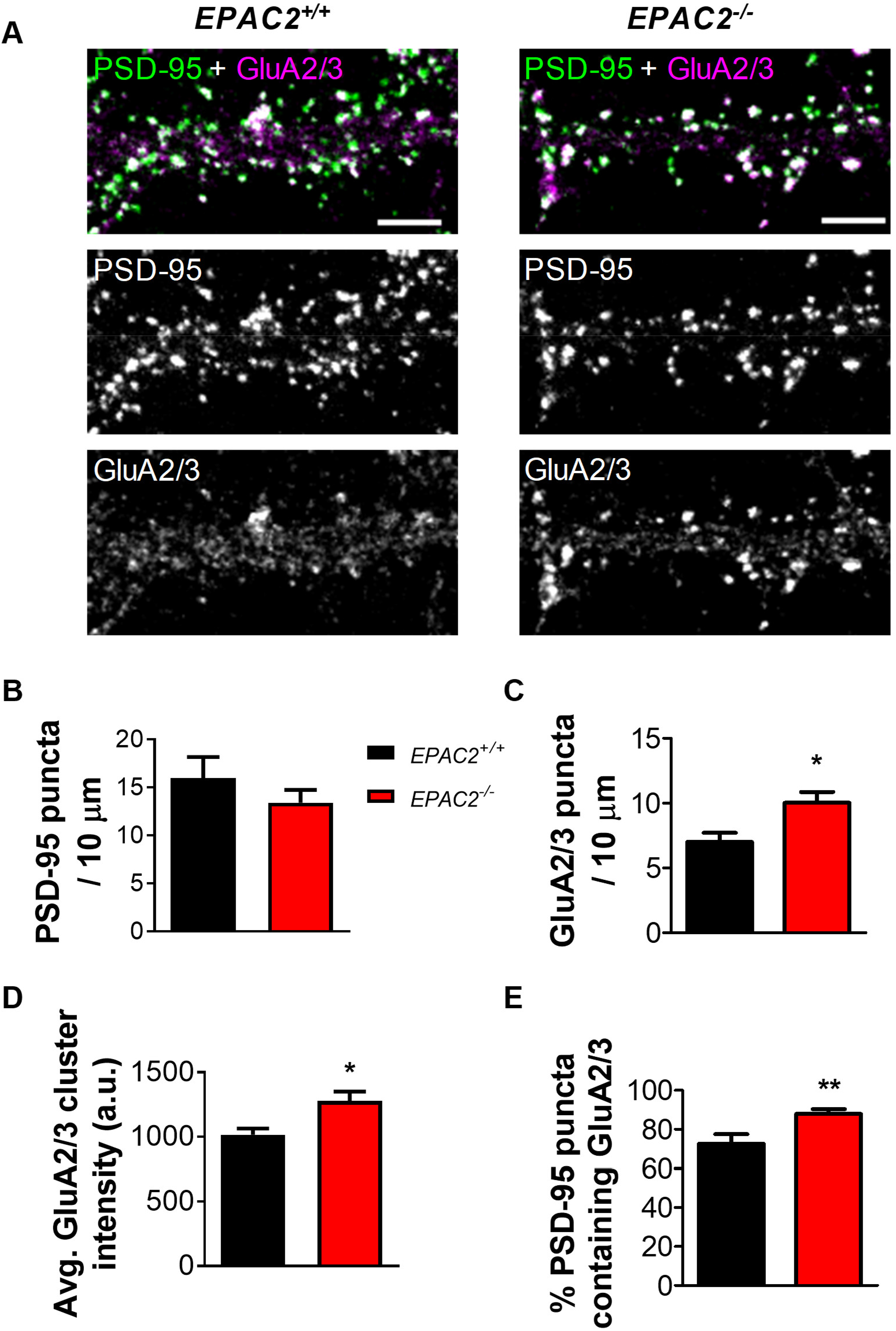
Loss of EPAC2 increases the colocalization of PSD-95 and GluA2/3 in dendrites. **(A)** Confocal images of cultured cortical neurons (DIV 22) from wild-type or *Epac2^−/−^* mice, immunostained for PSD-95 and GluA2/3. Scale bars=5 μm. **(B)** Quantification of linear density of PSD-95 puncta in neurons in panel A. **(C)** Quantification of GluA2/3 puncta density and comparison of genotypes reveals increased density of GluA2/3 puncta. *P=0.0277, Student’s t test; n=8-11 cells per condition, 3 independent cultures/genotype. **(D)** Quantification average cluster intensity and comparison of genotypes of GluA2/3 puncta. *P=0.0112; Student’s t test; n=8-11 cells per condition, 3 independent cultures/genotype. **(E)** Quantification of colocalization, as measured by percentage of PSD-95 puncta containing GluA2/3 immunofluorescence signal, in wild-type and *Epac2^−/−^* neurons. **P=0.0066, Student’s t test; n=8-11 cells per condition, 3 independent cultures/genotype. Data in the bar graphs are presented as mean ± s.e.m.

### Epac2^−/−^ neurons display altered adhesion protein expression at synapses

Loss of *Epac2* either *in vitro* or *in vivo* results in the destabilization of synapses, which is accompanied by an increase the presences of spines with larger spine heads (Srivastava et al., 2012a; Woolfrey et al., 2009). Synapse stability is coordinated by adhesion molecules such as N-cadherin and the neuroligins (Jang et al., 2017). We have previously shown that NL3 forms a protein complex with EPAC2 at synapses (Woolfrey et al., 2009). Moreover, Rap1 regulates the presence of N-cadherin at synapses (Xie et al., 2008). Therefore, we reasoned that, as *Epac2r^−/−^* cultures displayed abnormal dendritic spine morphologies and that EAPC2 is a direct regulator of Rap1, that neurons lacking this protein may also display altered expression of adhesion proteins at synapses.

We first examined the presence of NL3 and the pre-synaptic and active zone marker bassoon in wild-type and knockout cultures. Assessment of the linear density of bassoon revealed no difference between genotypes (**Figure 3A-B**). This is consistent with there being no alteration in synapse density in *Epac2r^−/−^* neurons.

**Figure 3.**
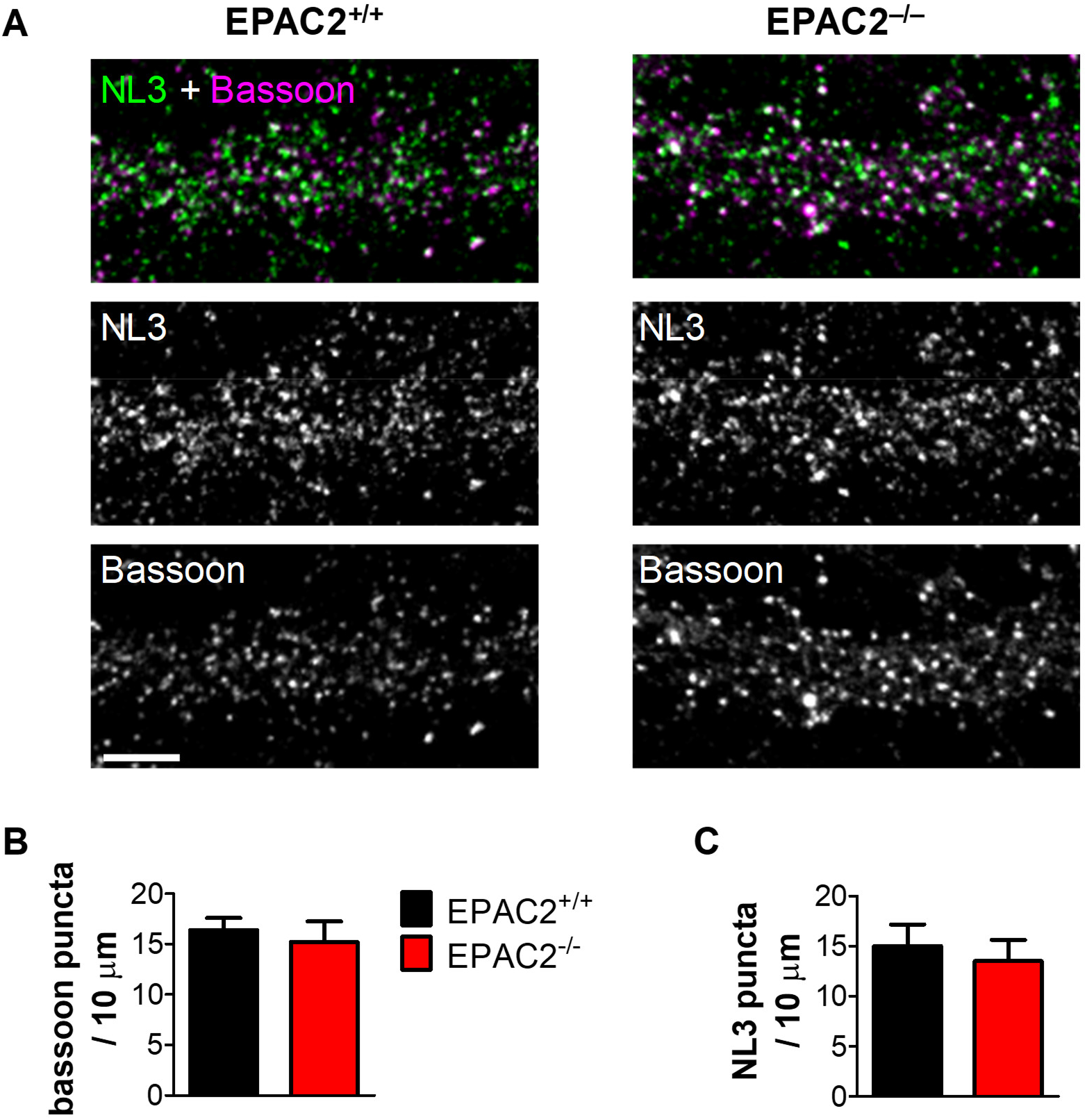
Loss of EPAC2 does not change puncta densities of neuroligin-3 or bassoon in dendrites. **(A)** Representative confocal images of cortical neurons (DIV 22) from wild-type (Epac2^+/+^) or *Epac2^−/−^* mice double immunostained for neuroligin-3 (NL3) and bassoon. Scale bar=5 μm. **(B-C)** Quantification of bassoon (B) and NL3 (C) puncta linear density in wild-type and *Epac2^−/−^* neurons. Genotypes were compared by Student’s t tests; n= 9-10 cells per condition, 3 independent cultures/genotype. Error bars represent s.e.m.

Interestingly, no difference in NL3 puncta density or size was observed between wild-type and *Epac2* knockout cultures. Furthermore, the number of NL3 and bassoon colocalizing puncta was similar in neurons from wild-type and *Epac2r^−/−^* cultures (**Figure 3A, C**).

N-cadherin is known to stabilize synapses and promote dendritic spine enlargement, an effect regulated by Rap1 activity (Xie et al., 2008). We therefore examined whether loss of EPAC2 would impact the clustering of N-cadherin at synapses. Again, we observed no difference in PSD-95 linear density between genotype (**Figure 4A and B**). Interestingly, we also observed no changes in the linear density of N-cadherin puncta between wildtype and knockout cultures (**Figure 4A and C**). However, when we examined N-cadherin puncta in more detail, we found that the cluster size was significantly increased in *Epac2^−/−^* cultures (**Figure 4A and D**). Critically, when we examined the density of colocalized PSD-95 and N-cadherin puncta, we found a significant increase in colocalized puncta in *Epac2r^−/−^* neurons, indicating an enrichment of this adhesion protein at synapses (**Figure 4A and E**). These data suggest that *Epac2r^−/−^* neurons have an increased amount of N-cadherin at synapses, consistent with an apparent increase in synapse stabilization.

**Figure 4.**
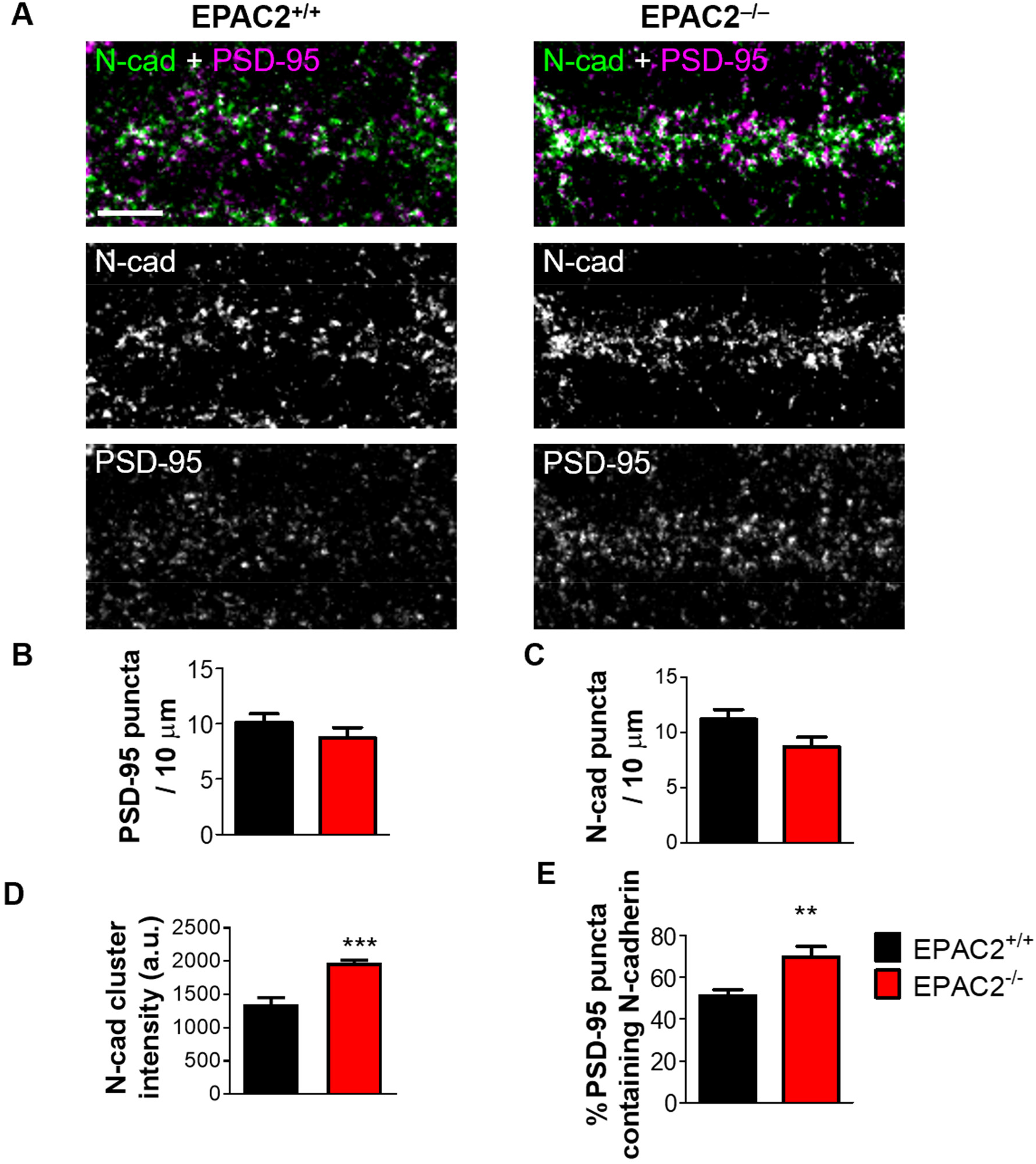
Loss of EPAC2 increases the colocalization of PSD-95 and N-cadherin in dendrites. **(A)** confocal images of cortical neurons from wild-type or *Epac2^−/−^* mice (DIV 22), immunostained for PSD-95 and N-cadherin (N-cad). Scale bar=5 μm. **(B-C)** Quantification and comparison of PSD-95 (B) and N-cadherin (C) puncta density between genotypes (Student t test). **(D)**. Quantification of N-cadherin puncta intensity revealed that average cluster size was larger in *Epac2^−/−^* neurons. ***P<0.001, Student’s t test; n=12-16 cells from 3 experiments/genotype. **(E)** Quantification of colocalization, as measured by the percentage of PSD-95 puncta that also contained N-cadherin immunofluorescence signal. Genotypes were compared by Student’s t test; **P=0.0056; n=12-16 cells from 3 experiments/genotype. Error bars represent s.e.m.

### Epac2^−/−^ neurons have altered balances of excitatory and inhibitory synapses

EPAC2 has been localized to both excitatory and inhibitory synapses (Woolfrey et al., 2009). Interestingly EPAC2 has been demonstrated to be important for excitatory transmission (Woolfrey et al., 2009; Yang et al., 2012) and has been shown to influence inhibitory transmission in dopamine neurons of the ventral tegmental area (Tong et al., 2017). Therefore, we were interested in examining whether loss of EPAC2 would impact excitatory and inhibitory synapses on the same neuron. Wild-type or *Epac2^−/−^* neurons (DIV 25) were immunostained with antibodies against vesicular glutamate transporter 1 (VGluT1) and vesicular GABA transporter (VGAT), presynaptic markers for excitatory and inhibitory synapses, respectively (**Figure 5A**). When we examined the linear density of these presynaptic markers along the dendrites of pyramidal neurons, we found a decrease in the ratio of VGluT to VGAT puncta in *Epac2r^−/−^* compared to wild-type neurons (**Figure 5B**). This effect appeared to be mediated not by any change in VGluT puncta density (**Figure 5C**), but rather a significant increase in VGAT puncta (**Figure 5D**). These data suggest that inhibitory drive is increased in the absence of EPAC2.

**Figure 5.**
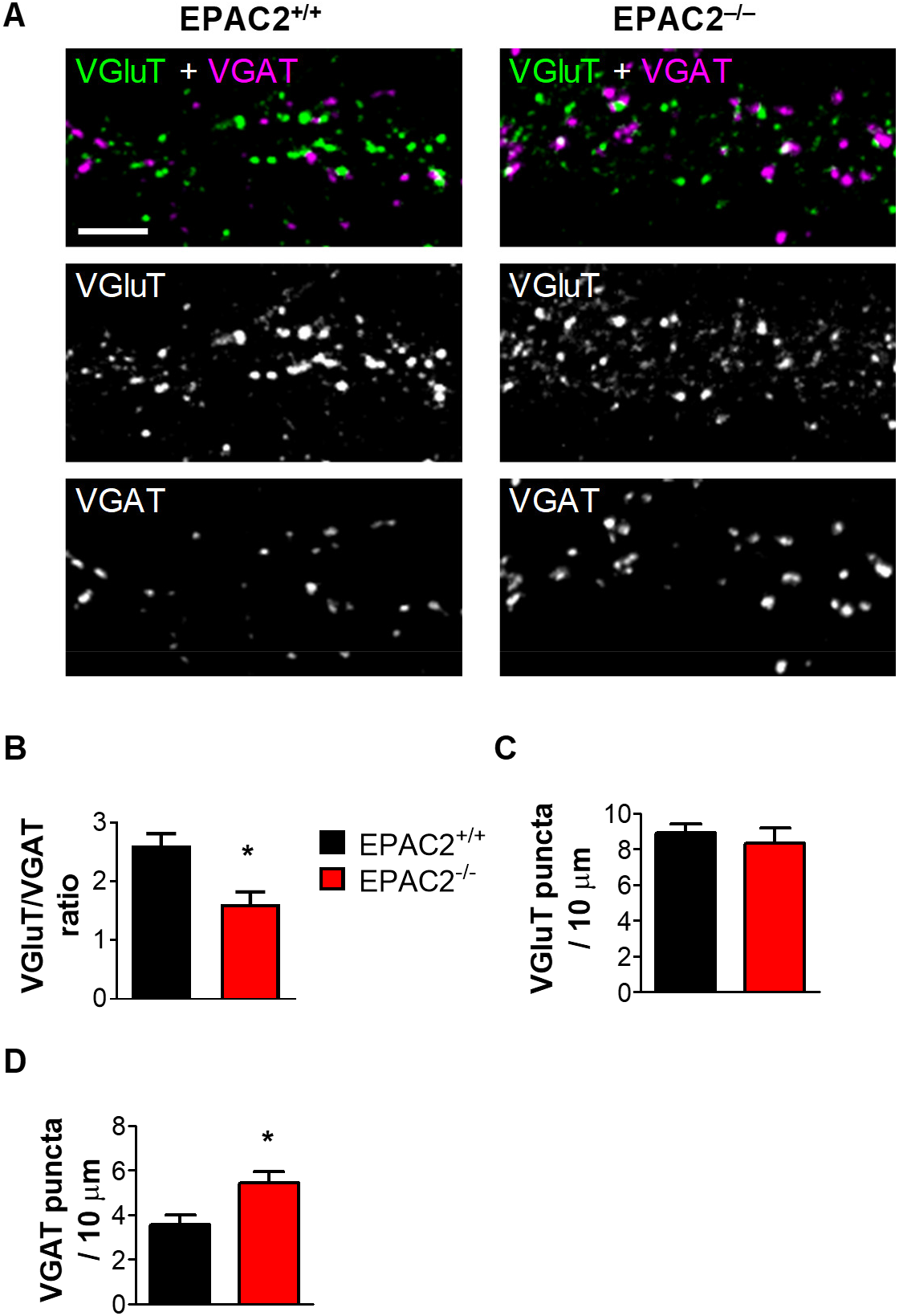
Loss of EPAC2 decreases the ratio of excitatory (VGlut) to inhibitory (VGAT) synaptic markers in dendrites. **(A)** Representative confocal images of cortical neurons from wild-type or *Epac2r^−/−^* mice (DIV 25), immunostained for VGluT1 and VGAT. Images were obtained by confocal microscopy. Scale bar=5 μm. **(B)** Quantification of the ratio of VGluT1 puncta density to VGAT puncta density. Genotypes were compared by Student’s t test, *P=0.0108; n=14-16 cells from 4 independent culture/genotype. **(C-D)** Quantification of VGluT1 puncta density. **(D)** Quantification of VGAT puncta density. Genotypes were compared by Student’s t test, *P=0.0276; n=14-16 cells from 4 independent culture/genotype. Error bars represent s.e.m.

### Epac2 is required for establishment of normal synaptic and dendritic structures in the ACC

We have previously shown that loss of *Epac2* alters the dendritic organization and spine dynamics of layer 2/3 and layer 5 neurons, respectively, located in premotor and somatosensory areas (Srivastava et al., 2012a; Srivastava et al., 2012b). Moreover, EPAC2 has been shown to be required for maintaining spine morphology as well as density *in vitro* and *in vivo* (Srivastava et al., 2012a; Woolfrey et al., 2009). As the analysis of EPAC2 effects on spine and dendritic morphologies had thus far been limited to the pre-motor and somatosensory areas, we were interested whether loss of *Epac2* also impacted these parameters in neurons from another cortical region. As *Epac2^−/−^* mice display gross disorganization of the ACC (Srivastava et al., 2012a), we therefore focused on the synaptic and dendritic morphologies of layer 5 neurons in this cortical region.

First, we assessed the linear density of spines on apical and basal dendrites of layer 5 neurons from the ACC of *Epac2^+/+GFP^* and *Epac2^-/-GFP^* mice. This analysis revealed no difference in the density of dendritic spines along apical or basal dendrites of layer 5 neurons in the ACC (**Figure 6A-B**). Analysis of spine morphology revealed that spine on apical dendrites of layer 5 ACC neurons in *Epac2^-/-GFP^* mice had significantly larger spine areas; there was no difference in spine area of spines on basal dendrites between wildtype and knockout animals (**Figure 6A-C**). These data provide further evidence that *Epac2* regulates spine stability in vivo.

**Figure 6.**
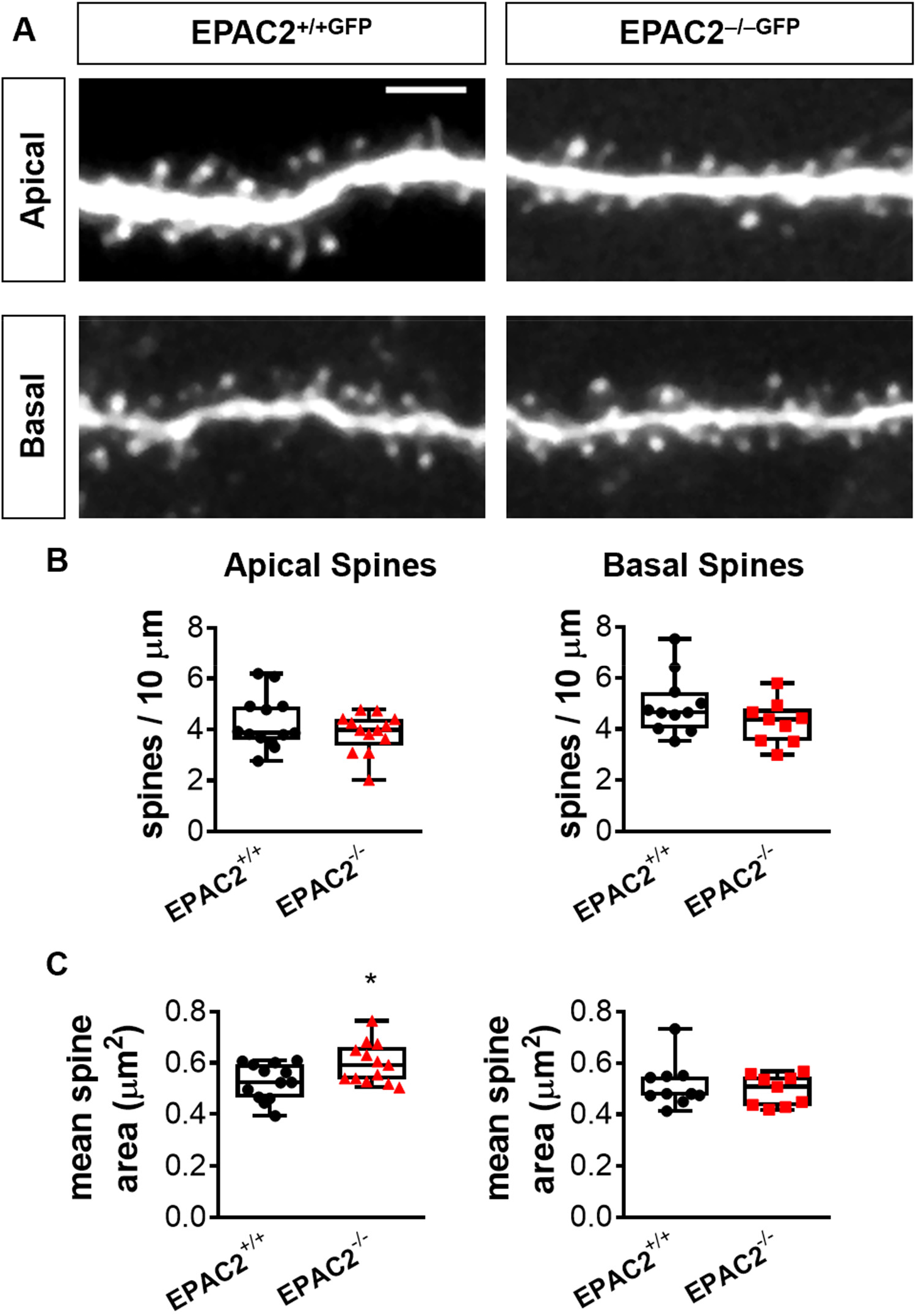
Dendritic spine morphology in layer 5 pyramidal neurons in the anterior cingulate cortex of Epac2-deficient mice. **(A)** Representative images of dendritic spines on apical or basal dendrites of layer 5 ACC neurons from *Epac2^+/+GFP^ or^-/-GFp^* mice. Images were acquired by 2PLSM. Scale bar=5 μm. **(B)** Quantification of spine linear density, of apical and basal dendritic spines in shown in panel A. Comparisons between genotypes were made by Student t test. **(C)** Quantification of dendritic spine area of apical and basal dendritic spines in ACC section prepared from Epac2^+/+GFP^ or *Epac2^-/-GFP^* mice. Comparisons between genotypes were made using Student’s t tests; *P=0.0214. Data were derived from 13 cells/genotype from 4 animals per genotype. Data are presented as means ± s.e.m.; each data point represents an individual cell.

Next, we examined the dendritic architecture of layer 5 ACC neurons in wildtype and *Epac2* knockout mice. As described for layer2/3 neurons in the somatosensory cortex (Srivastava et al., 2012b), layer 5 ACC neurons had a reduced number of basal, but not apical, dendrites (**Figure 7A and B**). Interestingly, both apical and basal dendrite branches were on average significantly longer in neurons from *Epac2^-/-GFP^* mice (**Figure 7C**). Consistent with these abnormalities, assessment of branch complexity as a function of branching order, revealed that both apical and basal higher order branch number were significantly reduced Epac2^-/-GFP^ mice versus wild-type mice (**Figure 7D**). These data are consistent with previous work demonstrating a role for EPAC2 in controlling the development of dendritic arborization.

**Figure 7.**
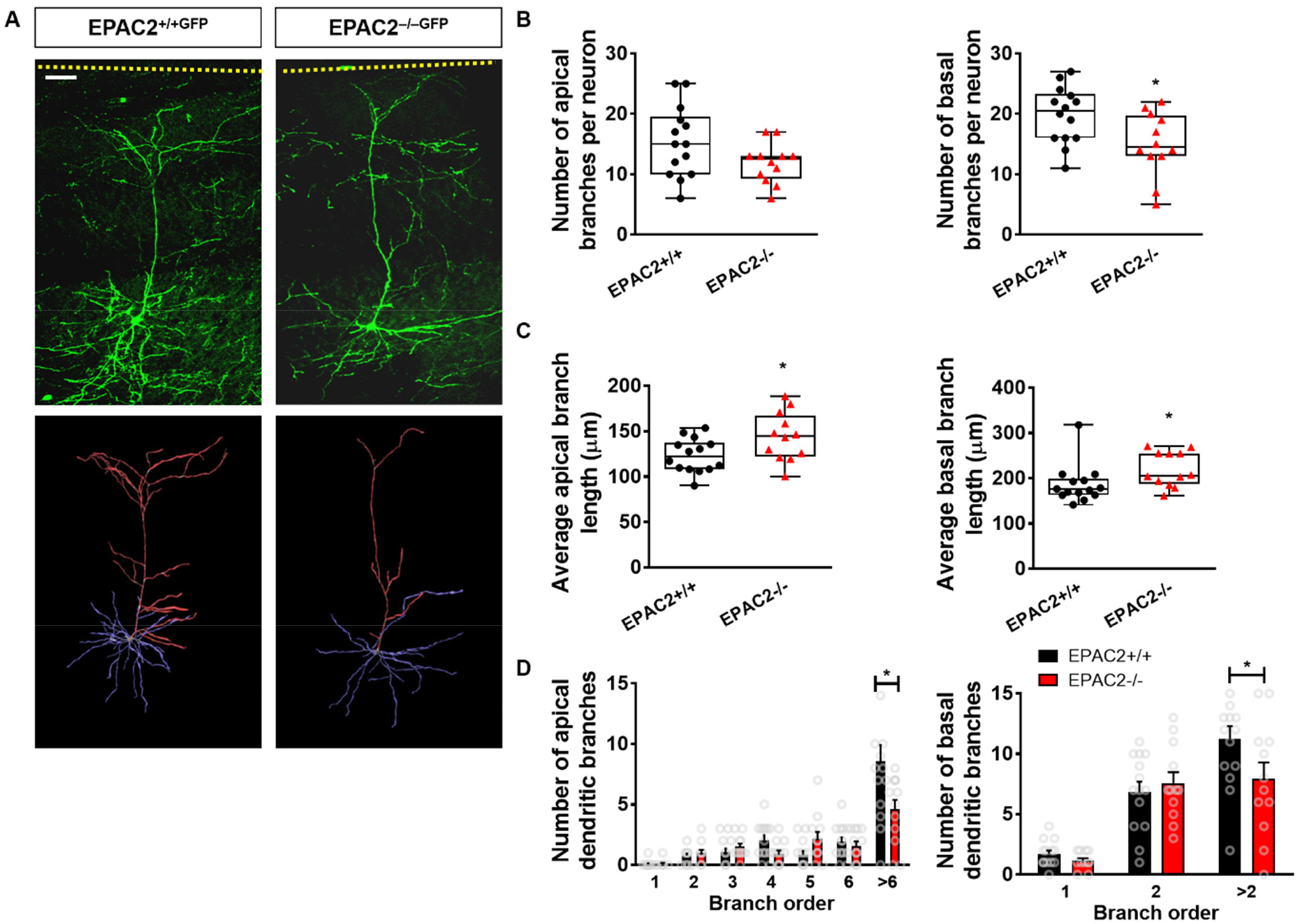
Epac2^-/-GFP^ mice display altered dendritic arborization of layer 5 ACC neurons. **(A)** Top: representative 2PLSM images of layer 5 pyramidal neurons in the ACC of 300-μm sections from *Epac2^+/+GFP^* and *Epac2^-/-GFP^* mice. Bottom: 3-D reconstructions of the apical (red) and basal (blue) dendritic arbors of neurons in the images above. Scale bar=50 μm. **(B)** Quantification of apical or basal dendritic branch number per neuron in layer 5 neurons in panel A. This revealed that there are fewer basal branches on layer 5 ACC neurons from *Epac2^-/-GFP^* mice. Comparisons between genotypes were made using Student’s t tests; *P=0.0216. Data were derived from 12-14 cells/genotype from 4 animals per genotype. **(C)** Assessment of average branch length for apical and basal dendrites of layer 5 pyramidal neurons in the ACC of 300-μm sections from *Epac2*^+/+GFP^ and *Epac2^-/-GFP^* mice. Apical and basal branches from *Epac2^-/-GFP^* mice were longer compared to wildtype mice. Comparisons between genotypes were made using Student’s t tests; *P=0.0265 (apical) or *P=.0478 (basal). Data were derived from 12-14 cells/genotype from 4 animals per genotype. **(D)** Quantification of apical or basal dendritic branching as a function of branch order in layer 5 neurons shown in panel A. The number of higher order dendritic branches on apical and basal dendrites were significantly decreased in *Epac2^−/−^* mice (mixed model ANOVA with Bonferroni post-tests; apical branch order: F(6, 168)=32.60, P<0.0001; genotype: F(1, 168)=3.223, P=0.0744; interaction: F(6, 168)=5.307, P<0.0001); basal branch order: F(2, 72)=45.42, P<0.0001; genotype: F(1, 72)=2.106, P=0.1511; interaction: F(2, 72)=2.746, P=0.0409). Data were derived from 12-14 cells/genotype from 4 animals per genotype. Data are presented as means ± s.e.m.; each data point represents an individual cell.

## Discussion

EPAC2 is a major PKA-indepodnet target for cAMP in the mammalian forebrain. Through its ability to regulate the small GTPase Rap1, EPAC2 is involved in regulating synapse stability (Woolfrey et al., 2009). In addition, EPAC2 is required for the establishment of basal dendritic arborization of layer 2/3 neurons in vivo (Srivastava et al., 2012b). Multiple studies have also shown that EPAC proteins are required for normal cognitive functions (Yang et al., 2012) with EPAC2 being required specifically for socio-communicative behaviors (Srivastava et al., 2012a) as well as playing a role in controlling anxious and depressive behaviors (Zhou et al., 2016). However, a comprehensive understanding of the role for EPAC2 in the development of the brain and organization of synapses is not fully understood. In this study, we confirm that EPAC2 is required for cAMP-mediated changes in spine morphology. Furthermore, we show that loss of EPAC2 results in the increased expression of GluA2/3-containing AMPA receptors and the adhesion protein N-cadherin at synapses. Interestingly, neurons from *Epac2^−/−^* also appear to have increased inhibitory input resulting in a potential imbalance in E/I ratio. Finally, we find that EPAC2 loss results in alterations in spine morphology and development of dendritic arborization of layer 5 ACC neurons in vivo. These data provide further evidence that correct expression of EPAC2 is required for the normal development of dendritic architecture, and moreover, is a critical regulator of synapse organization, particularly, in the establishment of synapse stability and E/I balance.

Analysis of cultured cortical neurons revealed that neurons from *Epac2^−/−^* mice had larger spine areas. This is consistent with previous work that has shown that EPAC2 is involved in regulating spine stability and motility (Srivastava et al., 2012a; Woolfrey et al., 2009). Larger spine would be more stable, have reduced motility and thus likely have altered responses to stimuli that would induced changes in spine morphology, ultimately impacting the ability of neural circuitry to response to plasticity inducing stimuli (Kasai et al., 2010; Penzes et al., 2011). Interestingly, we did not observe any change in the expression of NL3, a binding partner of EPAC2 (Woolfrey et al., 2009). This suggests that EPAC2 is not involved in regulating the expression of this adhesion protein but is only required for NL3-medaited signaling. However, concurrent with an increase in spine size, we observed an increase in the size of N-cadherin puncta at synapses in *Epac2* knockout cultures. An increase in N-cadherin at synapses has previously been shown to be linked with larger, more stable spines (Mendez et al., 2010; Xie et al., 2008). Thus, an increase in the amount of N-cadherin at synapses would be in line with larger spines more stable spines.

Previous work has shown that EPAC2 activation decreases surface expression of GluA2/3 and AMPA-mediated transmission (Woolfrey et al., 2009). Consistent with these results, we found that *Epac2r^−/−^* neurons had an increased density of GluA2/3-containing AMPA receptors, specifically at synapses. EPAC proteins and EPAC2 have been shown to be required for cAMP-dependent long-term depression (LTD) as well as cocaine induced switching of AMPA receptor subunit composition (Liu et al., 2016; Ster et al., 2009). The consequence of increased synaptic expression of GluA2/3-cotnaining AMPA receptors would potentially impact the ability of neurons to undergo plasticity-induced functional changes. Such deficits would also be consistent with our observation that loss of EPAC2 causes the formation of larger and more stable dendritic spines. Thus, EPAC2 appears to be required for maintaining the ability of neurons to undergo destabilization.

*Epac2r^−/−^* neurons also exhibited an increase in VGAT puncta, suggesting an increase of inhibitory synaptic input onto these neurons concurrent with increased synaptic glutamate receptor content. It may be somewhat surprising that an increase in GluA2/3 puncta is not accompanied by an increase in VGluT1 puncta density. But this result may be explained by the fact that PSD-95 puncta numbers are not changing, indicating that excitatory synapse numbers are similar in the presence and absence of EPAC2. Taken together, these data support a model in which the absence of EPAC2 leads to over-stabilized excitatory synapses, which also leads to an increase in the number of inhibitory inputs as a homeostatic response to the likely strong glutamatergic transmission occurring at these over-stabilized synapses.

An interesting observation in this study is that layer 5 neurons located in the ACC exhibit subtle changes in both dendritic and synaptic structures. Knockdown of EPAC2 *in vivo* results in the loss of dendritic spines on apical and basal dendrites of layer 2/3 neurons (Srivastava et al., 2012b). In contrast, no change in spine density was observed on layer 5 neurons in the ACC, but an increase in the number of spines with a larger area were found on apical dendrites. Similar to what we have previously reported following knockdown of EPAC2 on layer 2/3 neurons (Srivastava et al., 2012b), *Epac2^−/−^* mice had reduced basal dendrite number and complexity on layer 5 neurons in the ACC. Interestingly, we also observed subtle alteration in the length of dendritic branches on both apical and basal dendrites of these cells. It is likely that the more pronounced effect of EPAC2 loss on basal dendrites is due to the asymmetrical distribution of the EPAC2 protein throughout the dendritic tree.

The *Epac2* gene and its protein product have been implicated in a number of psychiatric disorders. *EPAC2* is located in the 2q31-q32 region, which was identified by several genome-wide linkage studies as an important autism susceptibility locus (Buxbaum et al., 2001; Shao et al., 2002). Microdeletion of the 2q31.1 region has been associated with intellectual disability and developmental delay (Williams et al., 2010). Recent studies also identified several copy number variants (CNVs) in this region in patients with autism, as well as enrichments of CNVs disrupting genes involved in GTPase/Ras signaling in autistic patients (Marshall et al., 2008; Pinto et al., 2010). Several rare mutations in the *EPAC2* gene have been identified in subjects with autism (Bacchelli et al., 2003), and interestingly, several of the mutations altered protein function, spine morphology and dendritic architecture (Srivastava et al., 2012b; Woolfrey et al., 2009). Because abnormal social and communication behavior is characteristic of a number of neurodevelopmental and neuropsychiatric disorders, and that *Epac2* knockout mice show impaired socio-communicative behaviors (Srivastava et al., 2012a; Yang et al., 2012), gaining a greater understanding of EPAC2 function *in vivo* may provide further insight into the pathogenesis of these diseases.

## Acknowledgments

This work was supported by NIH grants R01MH071316 and R01MH097216 to P.P.; and NRSA Ruth L. Kirschstein Award F31MH085362 to K. A. J.; and grants from the Medical Research Council, MR/L021064/1, Royal Society UK (Grant RG130856), and the Brain and Behavior Foundation (formally National Alliance for Research on Schizophrenia and Depression (NARSAD); Grant No. 25957), awarded to D.P.S.

